# A cytotoxicity dataset for data-driven design of ionizable lipid nanoparticles

**DOI:** 10.64898/2026.10.31.763372

**Authors:** Nathan Liu, Qikui Zhu, Zheng-Rong Lu

## Abstract

Ionizable lipid nanoparticles (LNPs) are widely used to deliver nucleic acid therapeutics, but the design of safe LNPs remains challenging due to the complex activity relationships among structure, composition, and physicochemical properties. Here, we present a curated dataset, the Lipid Nanoparticle Cytotoxicity Database (LNPCD), which aggregates 1,413 LNP cytotoxicity measurements across 1,288 unique ionizable lipids drawn from 11 high-throughput combinatorial studies. Lipid structures were generated computationally from the published libraries and manually verified against the original publications. Each record pairs a machine-readable SMILES representation of the ionizable lipid with its measured viability endpoint and the associated experimental descriptors, including cell line, dose, and cargo type. The dataset provides a traceable resource for examining how ionizable lipid structure and experimental conditions relate to in vitro cell viability, and for developing and evaluating computational models of lipid nanoparticle cytotoxicity.

## Background & Summary

Ionizable or protonatable lipid nanoparticles (LNPs) are a class of non-viral nucleic acid delivery systems that respond to pH changes in the subcellular compartments during the delivery process^1–3^. Ionizable LNPs demonstrated highly efficient cytosolic delivery and excellent safety profiles^4,5^. Ionizable lipid nanoparticles are clinically established delivery systems for nucleic acid therapeutics, such as Onpattro, Comirnaty, and Spikevax^6–9^. LNP formulations typically contain an ionizable lipid, a helper phospholipid, cholesterol, and a polyethylene glycol-conjugated lipid^10^. Among these components, the ionizable lipid has a central role in nucleic acid encapsulation, endosomal escape, delivery efficiency, and biocompatibility^11–15^. Consequently, extensive combinatorial libraries have been developed to identify ionizable lipids with improved delivery and tolerability profiles^16^.

Candidate LNPs are commonly evaluated using in vitro assays before more resource-intensive in vivo experiments, a process that becomes labor-intensive and costly at scale. The increasing size and chemical diversity of ionizable lipid libraries have therefore motivated the development of computational approaches for prioritizing candidate structures. In particular, machine learning (ML)-based models have been applied to reduce this screening burden by providing an in silico pre-screen before wet-lab testing^17–22^. However, their development remains constrained by the limited availability of high-quality training data. This limitation is particularly pronounced for toxicity, for which data are scattered across individual studies, reported using heterogeneous experimental protocols and outcome definitions, and have not yet been consolidated into a large, harmonized dataset suitable for systematic ML development.

Existing LNP resources primarily emphasize formulation composition, physicochemical characteristics, and delivery performance. LNPDB^23^ compiles LNP formulation and activity information and includes only a small number of hemolysis annotations (90 of approximately 20,000 records), whereas LNP Atlas^24^ provides standardized composition, physicochemical, and bioactivity information for published LNP formulations. These resources support broad analyses of formulation and delivery characteristics but do not systematically organize quantitative cell-viability measurements at the scale of combinatorial ionizable lipid libraries. A complementary resource centered on cell viability is therefore needed to support the analysis of associations among ionizable lipid structure, experimental context, and in vitro cytotoxicity.

Here, we present the Lipid Nanoparticle Cytotoxicity Database (LNPCD), a curated dataset of 1,413 cell-viability measurements associated with 1,288 ionizable lipids from 11 published high-throughput combinatorial studies. For each source library, ionizable lipid structures were reconstructed from the reported chemical components and reaction schemes, represented as canonical SMILES strings, and linked to the corresponding viability measurements. Source-reported values were retained alongside harmonized endpoints, and available experimental descriptors, including cell model, nucleic acid cargo, formulation composition, and dose, were recorded to preserve the context of each measurement.

Where existing LNP resources document delivery efficacy, LNPCD documents toxicity, so that efficacy and safety can be considered jointly in predictive modeling and design; similar curated, openly licensed collections that pair activity and toxicity endpoints have supported molecular property prediction in other areas of chemical and biomedical research^25,26^. The resulting resource provides traceable, machine-readable data for examining in vitro cytotoxicity patterns and for developing computational models while accounting for differences among source studies and experimental conditions.

## Methods

The overall curation pipeline is shown in Figure 1.

**Figure 1.**
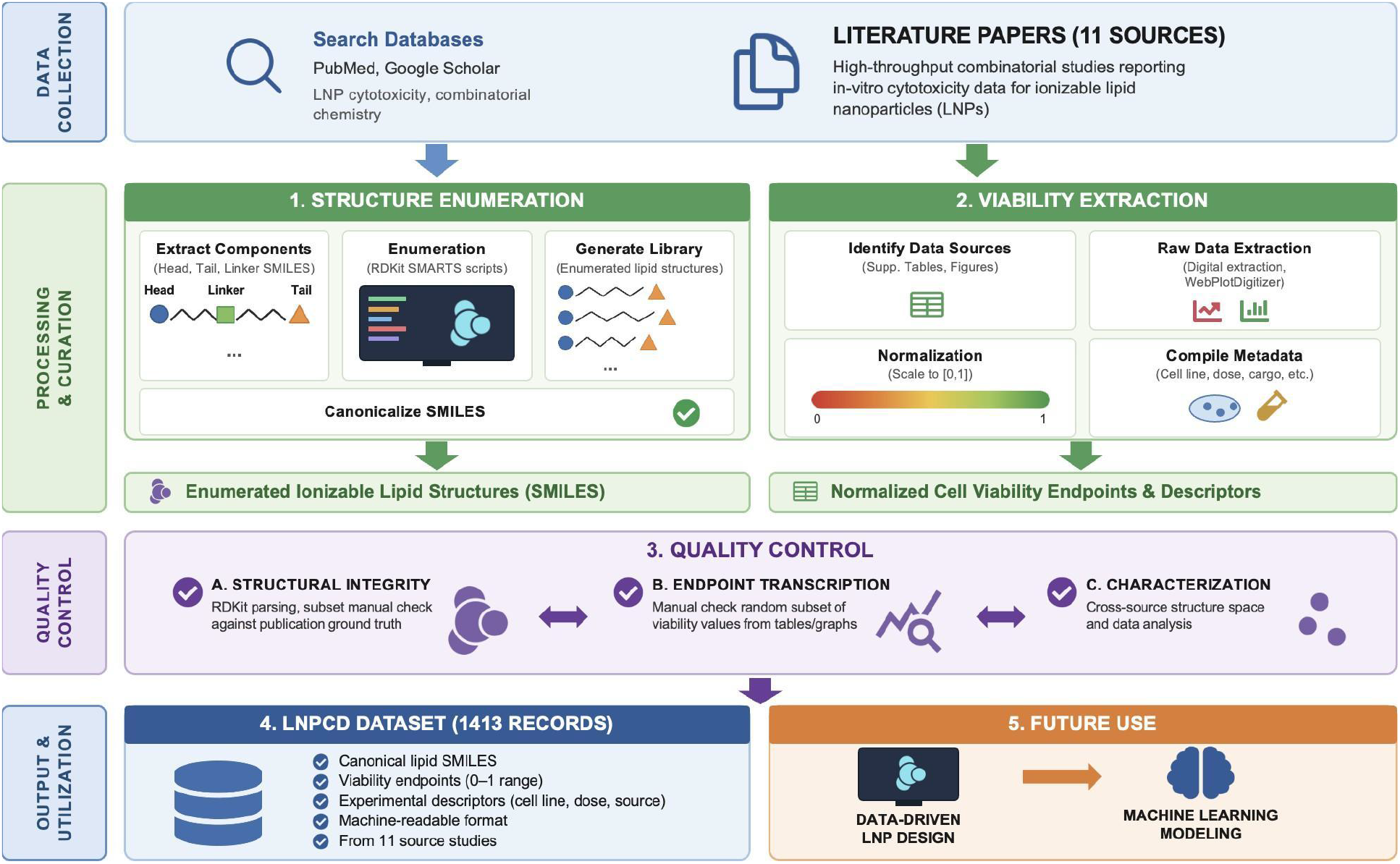
Construction and validation of the ionizable lipid nanoparticle cytotoxicity dataset. Source studies were identified and screened according to predefined eligibility criteria. Ionizable lipid structures were reconstructed from reported chemical components and reaction schemes and represented as canonical SMILES strings. Cell-viability values and associated experimental metadata were extracted from numerical source data or digitized from published figures, then converted into consistent data representations while retaining the source-reported values. Structural records, endpoints and metadata were evaluated using automated checks and manual auditing before release of the final dataset.

### Literature sourcing

We collected peer-reviewed publications reporting in vitro cell-viability measurements for ionizable lipid nanoparticle libraries. Searches were performed in PubMed and Google Scholar, combining LNP-related terms (“lipid nanoparticle,” “ionizable lipid,” “LNP”) with screening-related terms (“combinatorial library,” “high-throughput screening”) and safety-related terms (“cytotoxicity,” “cell viability,” “toxicity”). A study was included if it reported original cytotoxicity data for at least 25 unique LNP measurements, where a single ionizable lipid tested in multiple formulations or at multiple doses contributes one record per condition, and if the structural data for those formulations were fully recoverable. Studies that did not report formulation, structure, or viability details were excluded. Eleven high-throughput combinatorial chemistry studies remained and are listed in Table 1; the screening process is summarized in Figure 2.

**Table 1.** Source studies aggregated in the dataset. The source identifier used in the Experiment_ID column, the number of records contributed, and the number of unique lipids for each study.

| **Source ID** | **N (Records)** | **N (Unique Ionizable Lipids)** |
|---|---|---|
| Xue_CAD_LNP<sup>27</sup> | 180 | 180 |
| Lee_unsat<sup>28</sup> | 89 | 89 |
| Yu_aminoglycosides<sup>29</sup> | 44 | 44 |
| Farbiak_dendrimer<sup>30</sup> | 72 | 9 |
| Zhang_aminoglycosides<sup>31</sup> | 29 | 29 |
| Miller_zwitter<sup>32</sup> | 69 | 69 |
| Han_amidine<sup>33</sup> | 100 | 100 |
| Han_branched<sup>34</sup> | 93 | 31 |
| Han_a3<sup>35</sup> | 90 | 90 |
| Lin_peptide<sup>36</sup> | 75 | 75 |
| Liu_iphos<sup>37</sup> | 572 | 572 |
| **Total** | **1,413** | **1,288** |

**Figure 2.**
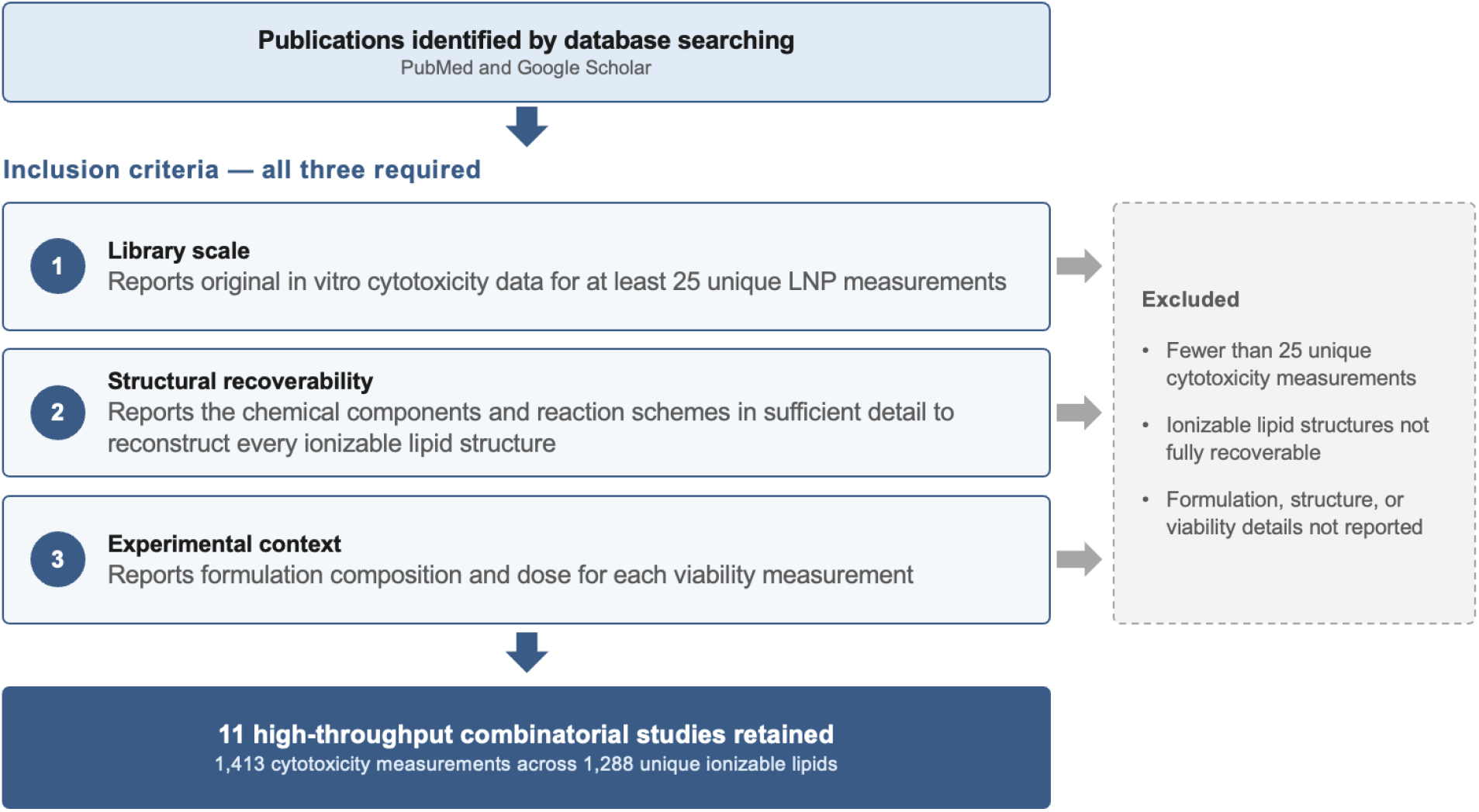
Study inclusion criteria. Publications were screened against three requirements: original in vitro cytotoxicity data for at least 25 unique LNP measurements, sufficient reporting of chemical components and reaction schemes to reconstruct the full ionizable lipid structures, and reported formulation and dose information for each measurement. Eleven high-throughput combinatorial studies, shown in Table 1, met all three and contributed the 1,413 measurements in LNPCD.

### Structure enumeration

Most source studies reported their combinatorial libraries by listing head group and tail group components, and linkers where applicable, rather than by drawing every full structure. The complete set of ionizable lipid structures can therefore be reconstructed computationally. Python scripts using RDKit^38^ SMARTS transformations combined the reported components according to the reaction chemistry described in each source. Generated SMILES^39^ were canonicalized with RDKit and then a random subset of 5-10% per study was checked by hand against the source paper for structural consistency. The source code used to generate these SMILES is available (see Code Availability).

### Derived structural features

We selected four structural features, number of unsaturated C-C bonds, number of protonatable nitrogens, number of carbons in the tail, and number of tails, that have been linked specifically to LNP activity and safety^11,15,40^. Structural descriptors were computed from the canonical SMILES with RDKit SMARTS patterns. Additionally, nine general molecular property features were selected and computed with RDKit. Every derived column is defined in Table 2.

**Table 2.** Column definitions.

| Column | Description |
|---|---|
| viability | Cell viability relative to untreated control, capped at 100% and divided by 100. |
| viability_raw_percent | Viability as reported in the source, in percent, before capping and scaling. |
| smiles | Canonical SMILES of the ionizable lipid, generated by RDKit. |
| Lipid_name | Name or identifier of the lipid as given in the source publication. |
| Experiment_ID | Source study identifier, matching the Source ID column of Table 1. |
| Ionizable_Lipid_Mol_Ratio | Molar percentage of ionizable lipid in the formulation. |
| Helper_Lipid_Mol_Ratio | Molar percentage of helper lipid. |
| Cholesterol_Mol_Ratio | Molar percentage of cholesterol. |
| PEG_Lipid_Mol_Ratio | Molar percentage of PEG lipid. |
| Helper_lipid_ID | Identity of the helper lipid used. |
| Ionizable_Lipid_to_mRNA_weight_ratio | Mass ratio of ionizable lipid to nucleic acid cargo. |
| Lipid_concentration | Ionizable lipid concentration in the well, in µg/mL. |
| NA_concentration | Nucleic acid concentration in the well, in µg/mL. |
| Lipid/Cells | Ionizable lipid mass per 1,000 cells, in ng. |
| NA/Cells | Nucleic acid mass per 1,000 cells, in ng. |
| Cargo_type | Nucleic acid delivered. Values: mRNA, siRNA. |
| Model_type | Cell line used in the cytotoxicity assay. |
| Num_tails | Number of tails on the ionizable lipid, where a tail is a string of carbons. |
| Num_carbon_in_tail | Number of carbons in the longest tail of the ionizable lipid. |
| MolWt | Molecular weight of the full ionizable lipid. |
| Topological_Polar_Surface_Area | TPSA of the ionizable lipid, in Å<sup>2</sup>. |
| Hydrogen_Bond_Donors | Number of HBD in the ionizable lipid. |
| LogP | Wildman–Crippen logP. |
| Fraction_sp3_Carbons | Fraction of carbon atoms that are sp3-hybridized. |
| Nitrogen_Count | Total N atoms in the ionizable lipid. |
| Rings | Total ring count in the ionizable lipid. |
| Aromatic_Rings | Aromatic ring count in the ionizable lipid. |
| has_ester | Indicator variable for the presence of ester linker. |
| num_unsaturated_cc_bonds | Number of unsaturated carbon-carbon bonds (C=C and C≡C) in the ionizable lipid. |
| num_protonatable_nitrogens | Number of protonatable nitrogens in the ionizable lipid. |
| paper_link | URL of the source publication for the record. |

### Data extraction and harmonization

Cell viability endpoints were recorded as percentage viability relative to an untreated control. Exact values were taken from a publication’s supplementary raw data where these were available. Where they were not, values were read off figures using WebPlotDigitizer^41^, applied to heat maps and bar charts.

Values above 100% were capped at 100%. Percentages were then divided by 100 to place them on the interval [0,1], which makes the endpoint on the scale expected of proportion-based ML models, useful for downstream modeling tasks. The uncapped, unscaled values were retained in the viability_raw_percent column.

Experimental descriptors (e.g. cell line, cargo) reported alongside each measurement were extracted from the source publication and retained as metadata columns so that users can stratify or filter the records. Dose descriptors were converted to a common basis. The columns Lipid_concentration and NA_concentration are the mass concentration in µg/mL of ionizable lipid and of nucleic acid in the well. The columns Lipid/Cells and NA/Cells are the mass in ng per 1,000 cells.

## Data Records

The dataset is deposited at Zenodo^42^ and is released under CC BY 4.0. The deposit contains lnpcd.csv, a single UTF-8 comma-separated file of 1,413 data rows.

One row is one cytotoxicity measurement: a single ionizable lipid, formulated and assayed under one set of conditions in one source study. Rows are not aggregated across replicates or across doses. Where a source reports the same lipid at more than one dose or in more than one cell line, each measurement appears as its own row. Table 2 defines every column.

## Data Overview

As summarized in Figure 3, most records in the dataset are non-cytotoxic, which follows the skew of the published literature it was drawn from. 106 records (7.5%) have viability below 80%, 118 (8.3%) fall between 80% and 90%, and 1,189 (84.1%) are above 90%.

**Figure 3.**
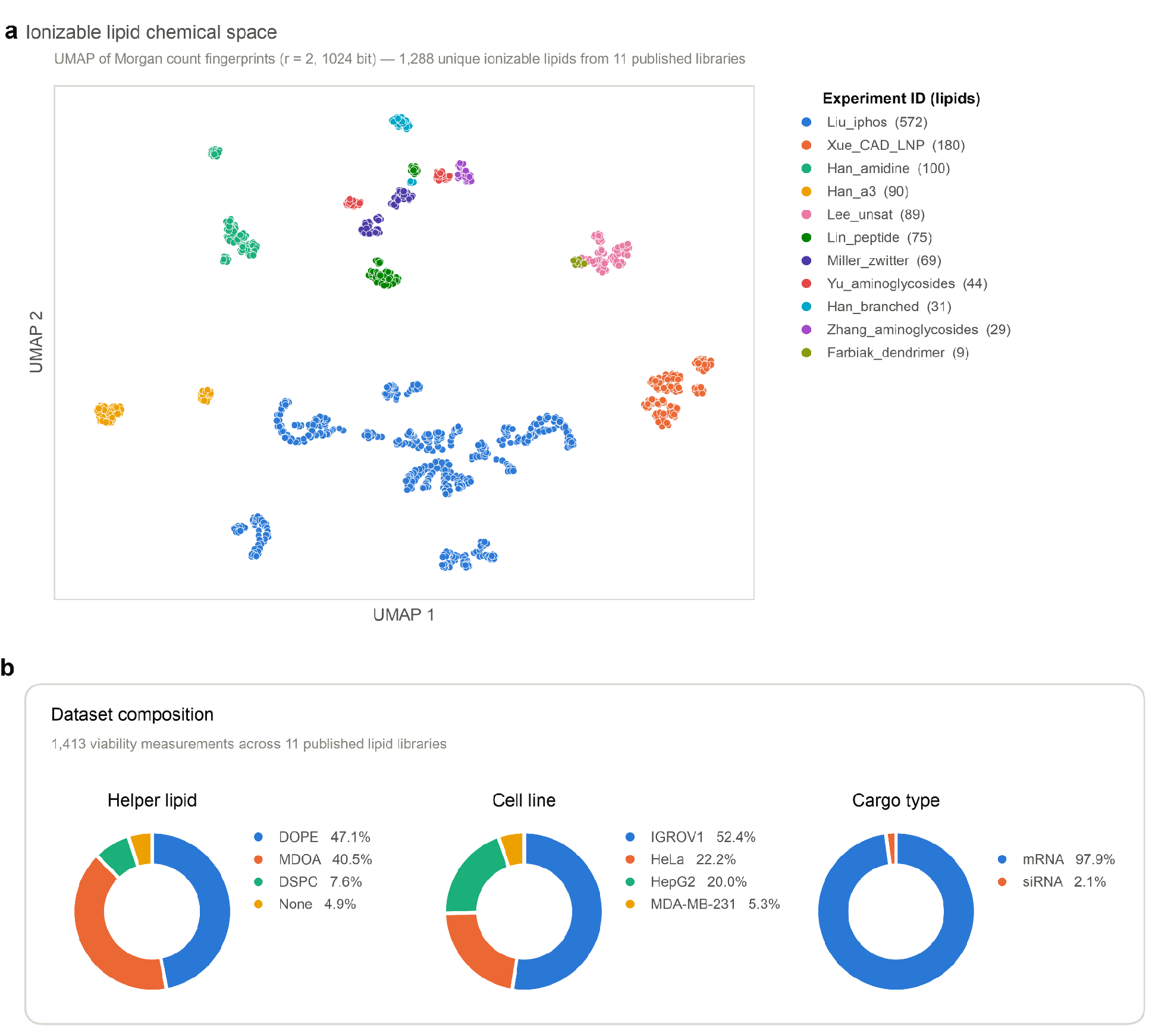
**a)** UMAP projection of the chemical space, colored by source library. Each point is one of the 1,288 unique ionizable lipids. Points were embedded from Morgan fingerprints^43^ of the SMILES, then projected to two dimensions with UMAP^44^. Axes are UMAP dimensions and carry no units. **b)** Statistics about the composition of LNPCD by helper lipid, cell line, and cargo type.

## Technical Validation

All 1,413 SMILES were parsed and canonicalized successfully with RDKit.

For each source study, a subset of approximately 5–10% of records was audited by hand against the source publication. In each audit the reviewer matched the SMILES structure against the structure reported in the source, confirmed the experimental conditions and derived features, and checked the viability endpoint against the original figure or table.

Automated checks were run over all 1,413 rows to catch out-of-range values or values that are impossible/highly unlikely, including viability in range, consistency between dosage features and weight ratio, SMILES valence, etc. The complete dataset was also screened for missing values, inconsistent category naming, and formatting errors.

## Usage Notes

Every record is machine-readable, pairing a SMILES string with tabular descriptors, and carries the identifier of the study it came from in Experiment_ID. Records can be filtered or stratified on that identifier, on cell line, on cargo, or on dose.

Two properties of the data affect how it should be modeled. First, because records are pooled from independent publications, chemical space and assay context are not evenly distributed across sources, and comparisons that cross source boundaries carry between-study noise^23,45^. Second, the viability distribution is heavily skewed toward non-toxic formulations, so sample weighting or a comparable correction is worth applying when training on the raw pooled table. The dataset also does not cover the full chemical and formulation space of ionizable lipids, so predictions on new chemotypes may amount to out-of-distribution extrapolation. Because all records are in vitro measurements, they should not be treated as proxies for in vivo tolerability; there is a known gap between in vitro and in vivo LNP behavior^46^.

## Data Availability

The dataset described here is available at Zenodo^42^ under DOI 10.5281/zenodo.23428876, released under CC BY 4.0. The deposit contains the data file lnpcd.csv. Source publications^27–37^ from which the records were extracted are given in Table 1.

A web application at https://lnpcd.pages.devrenders a searchable table of each datapoint in LNPCD. Users can click on each entry to see the SMILES as chemical diagrams and query records by lipid or experimental condition.

## Code Availability

Custom code used to enumerate the combinatorial libraries, compute the derived structural descriptors, and run the automated consistency checks is available at https://github.com/Nathan6006/LNPCD-SMILES/.

